# Hypothesis: Colony-forming activity of pluripotent stem cell-derived hepatocyte-like cells for stem cell assay

**DOI:** 10.1101/2021.11.30.470519

**Authors:** Kenta Ite, Masashi Toyoda, Saeko Akiyama, Shin Enosawa, Saeko Yoshioka, Takaaki Yukitake, Mayu Yamazaki-Inoue, Kuniko Tatsumi, Hidenori Akutsu, Hiroshi Nishina, Toru Kimura, Naoko Otani, Atsuko Nakazawa, Akinari Fukuda, Mureo Kasahara, Akihiro Umezawa

## Abstract

Hepatocyte-like cells (HLCs) generated from human pluripotent stem cells (PSCs) exhibit hepatocytic properties in vitro; however, their engraftment and functionality in vivo remain unsatisfactory. Despite optimization of differentiation protocols, HLCs did not engraft in a mouse model of liver injury. In contrast, organ-derived hepatocytes reproducibly formed colonies in the liver injury mouse model. As an extension of the phenomenon observed in hematopoietic stem cells giving rise to colonies within the spleen, commonly referred to as “colony-forming units in spleen (CFU-s“, we hypothesize that “colony-forming units in liver (CFU-L)“ serves as a reliable indicator of stemness, engraftment, and functionality of hepatocytes. The uniform expression of the randomly inactivated gene in a single colony, as reported by Sugahara et al. 2022, suggests that the colonies generated by isolated hepatocytes likely originate from a single cell. We, therefore, propose that CFU-L can be used to quantify the number of “hepatocytes that engraft and proliferate in vivo“ as a quantitative assay for stem cells that utilize colony-forming ability, similar to that observed in hematopoietic stem cells.

## INTRODUCTION

Hepatocyte-like cells (HLCs) derived from human pluripotent stem cells (PSCs) exhibit hepatocytic properties in vitro ^1–6^, albeit without engraftability in the urokinase-type plasminogen activator-cDNA/ severe combined immunodeficient (cDNA-uPA/SCID) mouse model. In contrast, hematopoietic stem cells (HSCs) exhibit colony-forming capacity, quantified as colony-forming units in spleen (CFU-s) ^7,8^. HSCs also display colony-forming ability in vitro (CFU-c: colony-forming units in culture) ^9^. HSCs generate many cell lineages from a single cell, and this clonality is an important feature of their function because it allows precise control of the blood system. Mesenchymal stromal cells also exhibit multipotency in vitro and in vivo ^10,11^. Likewise, we postulate that hepatocytes can be directly measured in vivo and a comparative analysis of HLCs derived from induced pluripotent stem cells (iPSCs) and organ-derived hepatocytes would enable the validation of the stemness, engraftability, and functionality of hepatocytes.

iPSCs have profoundly impacted areas of medicine, including regenerative therapy, drug discovery, and genetic disease research ^12–15^. Human iPSCs have also proved advantageous in toxicity studies. For instance, iPSC-derived hepatocytes have served as in vitro tools in drug metabolism ^16–19^. In this study, we used iPSCs derived from patients with drug-induced liver injury (DILI) because the immortality of iPSCs allows repeated acquisition of hepatocytes from the same origin.

Chimeric mice with humanized livers have been developed, among which cDNA-uPA/SCID mouse model shows a high replacement rate of human hepatocytes. This chimeric model by using isolated hepatocytes has proven to be useful in human drug metabolism and pharmacokinetics ^20–24^. To mimic normal human liver function, chimeric mice with humanized livers are usually generated by the transplantation of hepatocytes. Hepatocytes from patients with ornithine transcarbamylase deficiency (OTCD) form colonies in the liver of cDNA-uPA/SCID mice, and quite surprisingly, these colonies are presumably from a single hepatocyte ^25^. We here hypothesize that such colony-forming activity in liver (CFU-L) is directly related to the validation of stemness, viability, and functionality of hepatocytes. This hypothesis is an extension of the idea that bone marrow stem cells form colonies in the spleen ^7,8^.

## MATERIALS AND METHODS

### Ethical statement

Human cells in this study were obtained in full compliance with the Ethical Guidelines for Clinical Studies (Ministry of Health, Labor, and Welfare, Japan). The cells were deposited to RIKEN Cell Bank. Animal experiments were performed according to protocols approved by the Institutional Animal Care and Use Committee of the National Research Institute for Child Health and Development.

### Cells

Cells were obtained from a patient with DILI (Hep(C)). The cells were maintained in Dulbecco’s modified Eagle’s medium (DMEM, SIGMA D6429) supplemented with 20% FBS at 37°C in a humidified atmosphere containing 95% air and 5% CO_2_. When the cultures reached subconfluence, the cells were harvested with Trypsin-EDTA solution (cat# 23315, IBL CO., Ltd, Gunma, Japan), and re-plated at a density of 5 × 10^5^ cells in a 100-mm dish. Medium changes were carried out twice a week thereafter.

### Generation of iPSCs

iPSCs were generated from the patient-derived cells through a reprogramming by Sendai virus infection-mediated expression of OCT4/3, SOX2, KLF4, and c-MYC as previously described ^26^. Edom22iPS#S31 menstrual blood-derived cells were used for comparison purposes ^27,28^. Human iPSCs were maintained on irradiated mouse embryonic fibroblasts as previously described ^29,30^. The elimination of Sendai virus was confirmed by RT-PCR. Cells just after infection served as a positive control. Sequences of the primer sets for the Sendai virus are shown in Table 1.

**Table 1.**
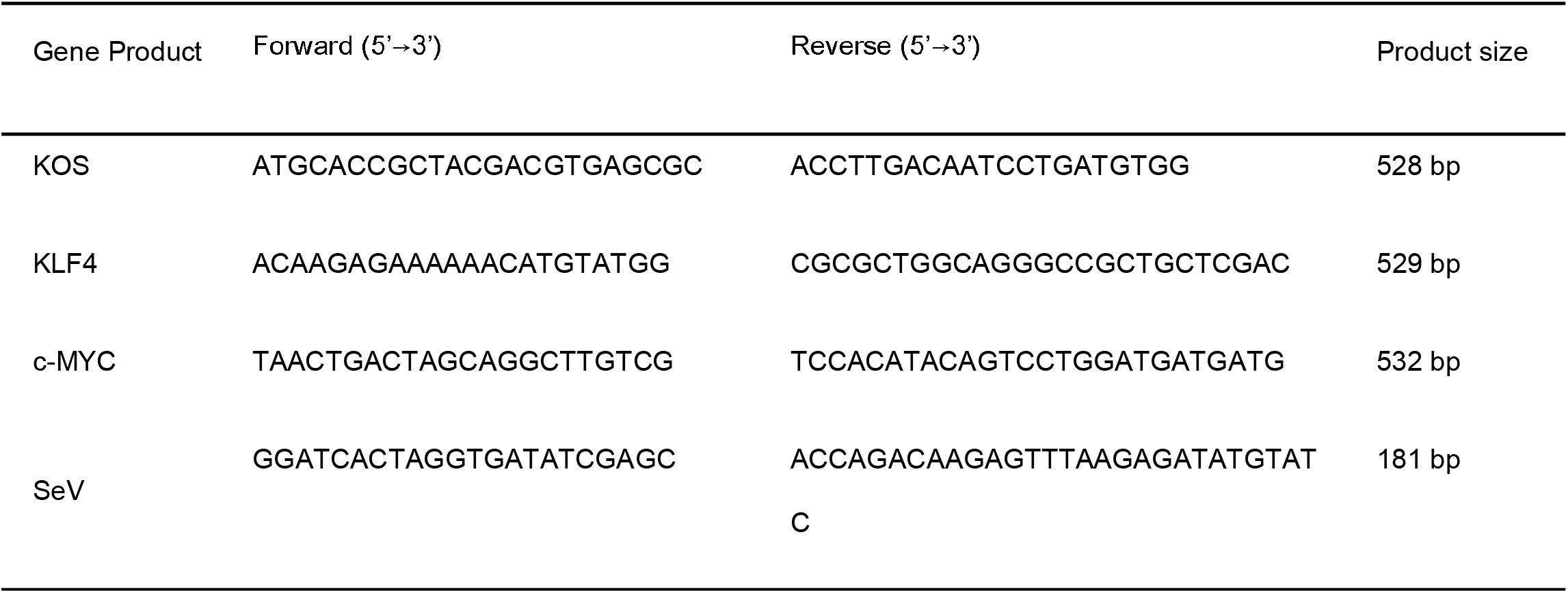
Primer sets for quantitative RT-PCR

### Immunocytochemical analysis

Cells were fixed with 4% paraformaldehyde in PBS for 10 min at 4°C. After washing with PBS and treatment with 0.2% Triton X-100 in PBS for 10 min, cells were pre-incubated with blocking buffer (10% goat serum in PBS) for 30 min at room temperature and then exposed to primary antibodies in blocking buffer overnight at 4°C. Following washing with 0.2% PBST, cells were incubated with secondary antibodies; either anti-rabbit or anti-mouse IgG conjugated with Alexa 488 or 546 (1:300) (Invitrogen) in blocking buffer for 30 min at room temperature. Then, the cells were counterstained with DAPI and mounted.

### Karyotypic analysis

Karyotypic analysis was contracted out to Nihon Gene Research Laboratories Inc. (Sendai, Japan). Metaphase spreads were prepared from cells treated with 100 ng/mL of Colcemid (Karyo Max, Gibco Co. BRL) for 6 h. The cells were fixed with methanol:glacial acetic acid (2:5) three times and placed onto glass slides (Nihon Gene Research Laboratories Inc.). Chromosome spreads were Giemsa banded and photographed. A minimum of 10 metaphase spreads were analyzed for each sample, and karyotyped using a chromosome imaging analyzer system (Applied Spectral Imaging, Carlsbad, CA).

### Teratoma formation

iPSCs were harvested by Accutase treatment, collected into tubes, and centrifuged. The cell pellets were suspended in the iPSellon medium. The same volume of Basement Membrane Matrix (354234, BD Biosciences) was added to the cell suspension. The cells (> 1 × 10^7^) were subcutaneously inoculated into immunodeficient, non-obese diabetic (NOD)/severe combined immunodeficiency (SCID) mice (CREA, Tokyo, Japan). The resulting tumors were dissected and fixed with PBS containing 4% paraformaldehyde. Paraffin-embedded tissue was sliced and stained with hematoxylin and eosin. The operation protocols were approved by the Laboratory Animal Care and the Use Committee of National Center for Child and Health Development, Tokyo.

### Hepatic differentiation

To generate embryoid bodies (EBs), iPSCs (1 × 10^4/well) were dissociated into single cells with accutase (Thermo Scientific, MA, USA) after exposure to the rock inhibitor (Y-27632: A11105-01, Wako, Japan), and cultivated in the 96-well plates in the EB medium [76% Knockout DMEM, 20% Knockout Serum Replacement (Life Technologies, CA, USA), 2 mM GlutaMAX-I, 0.1 mM NEAA, Pen-Strep, and 50 μg/mL l-ascorbic acid 2-phosphate (Sigma-Aldrich, St. Louis, MO, USA)] for 10 days. The EBs were transferred to the 24-well plates coated with collagen type I and cultivated in the XF32 medium [85% Knockout DMEM, 15% Knockout Serum Replacement XF CTS (XF-KSR; Life Technologies), 2 mM GlutaMAX-I, 0.1 mM NEAA, Pen-Strep, 50 μg/mL L-ascorbic acid 2-phosphate (Sigma-Aldrich, St. Louis, MO, USA), 10 ng/mL heregulin-1β (recombinant human NRG-beta 1/HRG-beta 1 EGF domain; R&D Systems, Minneapolis, MN, USA), 200 ng/mL recombinant human IGF-1 (LONG R^3^-IGF-1; Sigma-Aldrich), and 20 ng/mL human bFGF (Life Technologies)] for 14 to 35 days.

### Implantation into immunodeficient mice

HepaKI cells were harvested by treatment with Liberase, collected into tubes, centrifuged, and resuspended in PBS (200 μl). The same volume (200 μl) of Matrigel was added to the cell suspension. The cells (> 1 × 10^7) were inoculated into the subcapsular space of the kidney in SCID mice (CREA, Tokyo, Japan). The resulting tumors were dissected and fixed with PBS containing 4% paraformaldehyde. Paraffin-embedded tissue was sliced and stained with HE. The operation protocols were accepted by the Laboratory Animal Care and the Use Committee of National Center for Child and Health Development, Tokyo. For in vivo engraftment in livers, PSC-derived hepatocytes and organ-derived hepatocytes were implanted into the spleens of 2 to 4-week-old, homozygous cDNA-uPA/SCID mice as previously described ^25,31^.

### Immunohistochemistry

Subcutaneous nodules on mouse backs were fixed in 20% formalin and embedded in paraffin. Cut paraffin sections were deparaffinized, dehydrated, and treated with 2% proteinase K (Dako Omnis) in Tris-HCl buffer solution (pH 7.5) for 5 min at room temperature, or heated in ChemMate Target Retried Solution (Dako Omnis) for 5-20 min in a high-pressure steam sterilizer for epitope unmasking. After washing with distilled water, samples were placed in 1% hydrogen peroxide/methanol for 15 min to block endogenous peroxidase. The sections were then incubated at room temperature for 60 min in primary antibodies diluted with antibody diluent (Dako Omnis). The following primary antibodies against various human differentiation antigens were used: CYP2C (sc-53245, Santa Cruz), vimentin (V9, M0725, Dako Omnis, Glostrup, Denmark), albumin (ALB) (Dako Omnis), and AE1/AE3 (712811, NICHIREI). Then, they were washed three times with 0.01 M Tris-buffered saline (TBS) solution (pH 7.4) and incubated with goat anti-mouse or anti-rabbit immunoglobulin labeled with dextran molecules and horseradish peroxidase (EnVision, Dako Omnis) at room temperature for 30 min. After three times washes with TBS, they were incubated in 3,3’-diaminobenzidine in substrate-chromogen solution (Dako Omnis) for 5-10 min. Negative controls were performed by omitting the primary antibody. The sections were counterstained with hematoxylin.

### Cytochrome P450 induction

The expression levels of three major CYP enzymes, CYP1A2, CYP2B6, and CYP3A4, were examined to evaluate CYP induction in HepaKI. HepaKI was treated with omeprazole, phenobarbital, and rifampicin, and controls were treated with dimethyl sulfoxide (DMSO), as described in the previous study^17^.

### Statistical analysis

Statistical analysis was performed using the unpaired two-tailed Student’s t-test.

## RESULTS

### Generation of drug-induced hepatic injury iPSCs

We generated iPSCs from a patient with hepatic failure by Sendai virus infection-mediated expression of OCT4/3, SOX2, KLF4, and c-MYC (Figure 1A). When these reprogramming factors were introduced into 2.0 × 10^5^ cells (Hep(c) cell) from this patient, 11 clones of iPSCs were successfully generated and designated as iPSC-K (clones #12, #19, #24, #25, #41, #47, #53, #62, #66, #100, #116; Figure 1B, C). The efficiency of the iPSC colony generation was low compared to that of undamaged human cells from various adult tissues. Morphological characteristics of iPSC-K, *i*.*e*. flat and aggregated colonies, were similar to those of normal iPSCs and embryonic stem cells (Figure 1C). RT-PCR analysis revealed the elimination of the Sendai virus (Figure 1D, E). Immunocytochemical analyses demonstrated expression of the pluripotent cell-specific markers, *i*.*e*. SSEA-4, TRA-1-60, SOX2, NANOG, and OCT4/3, which was consistent with the profile observed in iPSCs (Figure 1F).

**Figure 1.**
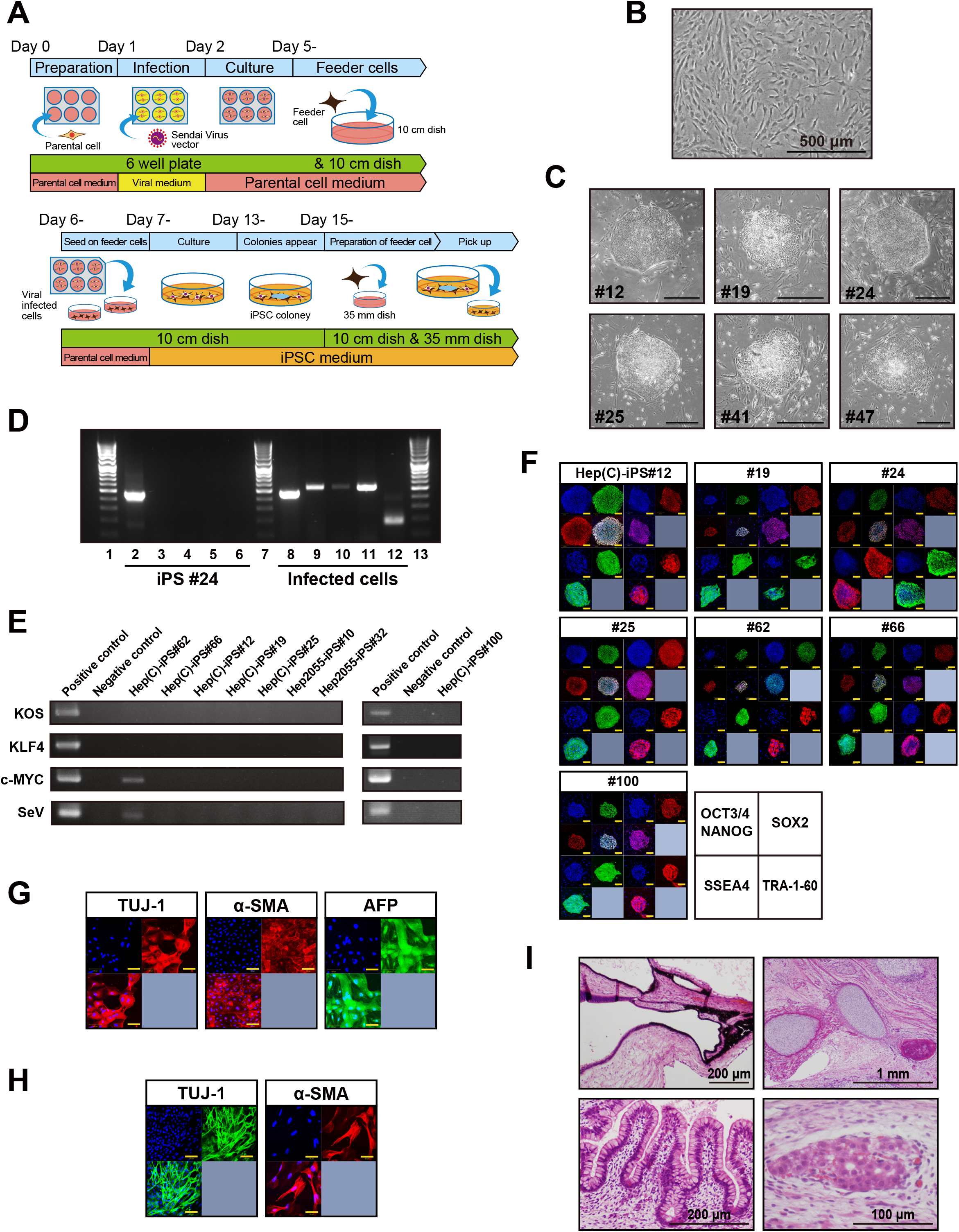
Generation of iPSCs from a patient with DILI. A. Scheme of iPSC generation. B. Phase contrast photomicrography of the parental cells (patient fibroblasts from the liver). C. Generation of iPSC clones (#12, #19, #24, #25, #41, #47). D. Elimination of Sendai virus vectors in iPSC-K#24. E. Elimination of Sendai virus vectors in the other clones. F. Immunocytochemistry of the pluripotency-associated markers, i.e. OCT4/3, NANOG, SOX2, SSEA4 and TRA-1-60, in iPSC-K (#12, #19, #24, #25, #62, #66, #100). G. in vitro differentiation of three germ layers of iPSC-K#24 via EB formation and adhesion culture in the EB medium. The differentiated cells were applied to the immunocytochemical analyses for markers of ectoderm (TUJ1), mesoderm (αSMA), and endoderm (AFP). H. Neural and muscular differentiation of iPSC-K#25 via EB formation and adhesion culture in the EB medium. The differentiated cells were applied to the immunocytochemical analyses for markers of ectoderm (TUJ1) and mesoderm (αSMA). I. Teratoma formation after subcutaneous injection of iPSC-K#25. iPSC-K#25 exhibited in vitro differentiation of three germ layers such as retinal pigmented epithelium (ectoderm), cartilage (mesoderm), intestinal epithelium (endoderm), and hepatocytes (endoderm).

To investigate multipotency in vitro, iPSCs were differentiated into ectodermal, mesodermal, and endodermal lineages. The differentiation of iPSC-K was confirmed by immunostaining using antibodies against β-tubulin III (TUJ1), α-smooth muscle actin (SMA), and α-fetoprotein (AFP) as ectodermal, mesodermal, and endodermal markers, respectively (Figure 1G, H). To address whether the iPSC-K has the competence to differentiate into specific tissues in vivo, teratomas were formed by implantation of iPSC-K in the subcutaneous tissue of immunodeficient NOD/SCID mice. iPSC-K produced teratomas within 6-10 weeks after implantation. Histological analysis of paraffin-embedded sections demonstrated that the three primary germ layers were generated as shown by the presence of ectodermal, mesodermal, and endodermal tissues in the teratoma (Figure 1I), implying iPSC-K has the potential for multilineage differentiation *in vitro* and *in vivo*. Among iPSC-K clones, #25, #66, and #100 generated a larger area of liver-like tissues in the teratomas, while the other clones did not.

### Hepatic differentiation

We investigated the efficiency of hepatic differentiation of iPSC-K by two different methods Protocol H and Protocol S (described in detail in Figure 2A, B, C). The differentiated cells exhibited hepatocyte-like morphology, i.e. a polygonal and/or cuboidal shape that had tight cell-cell contact when generated by either method (Figure 2D, E) and the expression of the hepatocyte-associated genes were comparable. Quantitative analysis revealed that iPSC-K expressed the genes for AFP, ALB, and α-antitrypsin (AAT) 21 days after the start of induction (Figure 2F, G, H). We employed Protocol S and used iPSC-K#25 hereafter for the iPSC-K experiments because iPSC-K#25 exhibited morphology that most resembled primary hepatocytes, and high expression of hepatocyte-associated genes. Time-course analysis revealed that similar expression levels of liver-associated genes were observed at 21, 28, and 35 days after the start of hepatic induction (Figure 2I, J).

**Figure 2.**
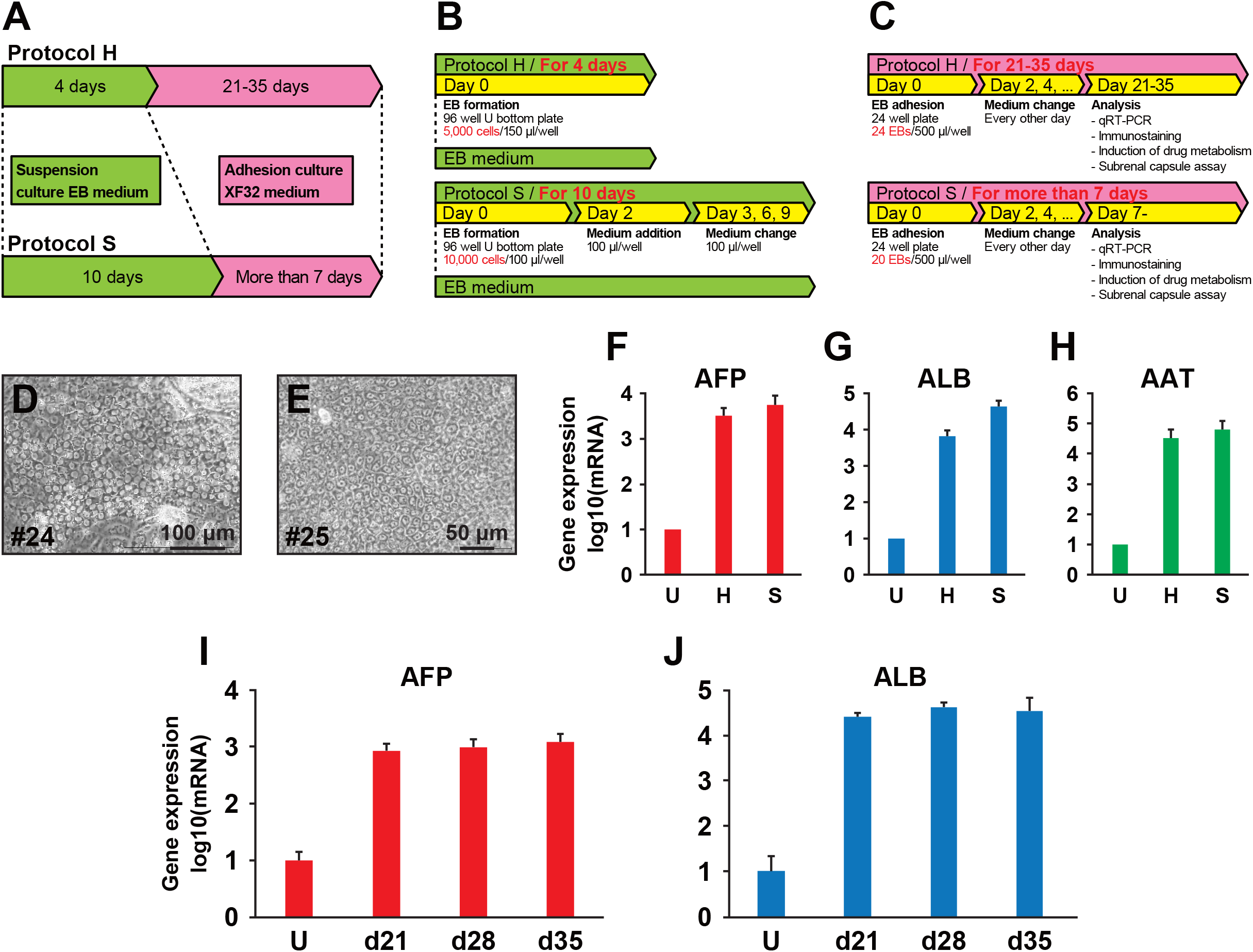
Protocols for hepatic differentiation of iPSCs. A. Scheme of two protocols, H and S. The protocols differ in the periods of the suspension and adherent culture for hepatic differentiation. B. Growth medium used for the suspension culture of Protocol H and S. C. Growth medium used for the adhesion culture of Protocol H and S. D. Phase-contrast photomicrograph of the hepatocyte-like cells (HLCs) generated from iPSC-K #24 with Protocol S. E. Phase-contrast photomicrograph of the HLCs generated from iPSC-K #25 with Protocol S. F. Expression of the α-fetoprotein (AFP) gene in the iPSC-K#25-derived hepatocytes at Day 21 with Protocol H and S. U: Undifferentiated iPSCs, H: Protocol H, S: Protocol S. G. Expression of the albumin (ALB) gene in the iPSC-K#25-derived hepatocytes at Day 21 with Protocol H and S. U: Undifferentiated iPSCs, H: Protocol H, S: Protocol S. H. Expression of the α-antitrypsin (AAT) gene in the iPSC-K#25-derived hepatocytes at Day 21 with Protocol H and S. U: Undifferentiated iPSCs, H: Protocol H, S: Protocol S. I. Time-course of the AFP gene expression with Protocol S in iPSC-K#25. RNAs were isolated at 21 days (d21), 28 days (d28), and 35 days (d35) after start of the EB formation. U: undifferentiated iPSCs without EB formation. J. Time-course of the ALB gene expression with Protocol S in iPSC-K#25. RNAs were isolated at 21 days (d21), 28 days (d28), and 35 days (d35) after start of the EB formation.

### Verification of the manufacturing process

To ensure the consistent production of HLCs from iPSC-K, we established a master cell bank and used a working cell bank (WCB) for starting material. We then developed a standard operating procedure for hepatic differentiation (Figure 3A). HepaKI developed into a cell type that was positive for both AFP and ALB (Figure 3B). Karyotypic analysis showed that the same chromosomes, without any aberration, were present in the WCB stock as parental fibroblastic cells from the patient (Figure 3C, D). Exome analysis revealed that iPSC-K had no significant single nucleotide alterations in a homozygous manner. To verify the procedure, we repeatedly manufactured HLCs from the WCB. Immunocytochemistry confirmed that these HLCs were positive for cytokeratin 7 (CK7), CK8/18 (AE1/3), and Hep1, but negative for CD31 and CD34 (Figure 3E).

**Figure 3.**
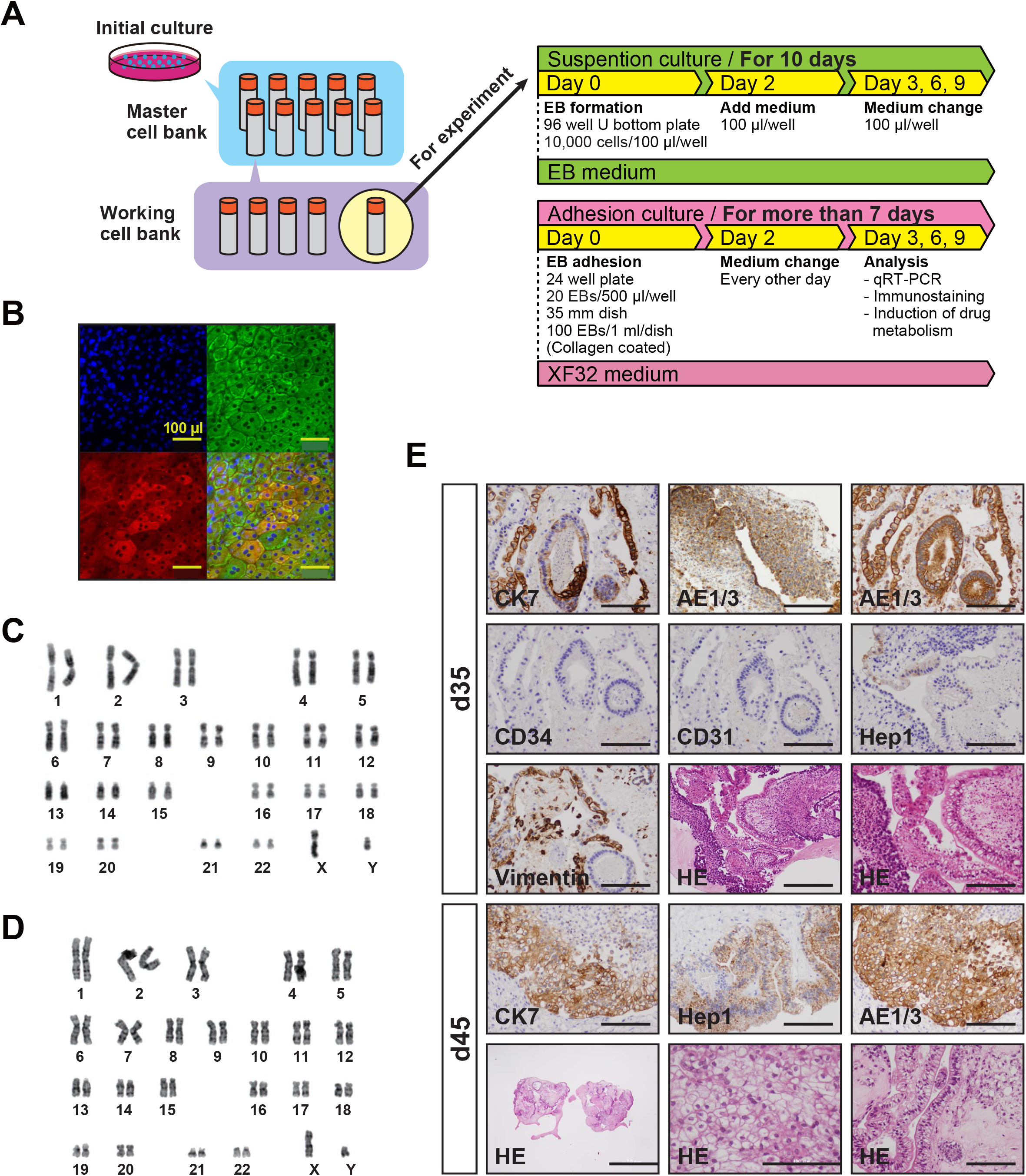
Systemization of drug-mediated CYP induction test. A. Scheme of the operation procedure for the CYP induction test with HepaKI cells. B. Immunocytochemistry of HepaKI with the antibodies to ALB (red) and AFP (green). C. Karyotypic analysis of parental fibroblastic cells from the patient’s liver. D. Karyotypic analysis of iPSC-K#25. E. Immunocytochemistry of HepaKI cells at 35 and 45 days in iPGel with the antibodies to cytokeratin 7 (CK7), cytokeratins 8/18 (AE1/3), CD34, CD31, CPS1 (Hep1), and vimentin. Bar: 100 μm. HE: hematoxylin and eosin stain.

### Induction of the genes for cyp1A2, 2B6, and 3A4

To investigate whether HepaKI exhibits CYP induction, we exposed HepaKI to omeprazole, phenobarbital, and rifampicin for 24, 48, and 48 h, respectively (Figure 4). Expression of the genes for AFP and ALB was unchanged with exposure to these drugs. Cyp1A2 was increased after exposure to omeprazole and phenobarbital, while Cyp2B6 remained unchanged (Figure 4C, D). Interestingly, Cyp3A4 was up-regulated 57.2-fold, on average, upon exposure to rifampicin (Figure 4E).

**Figure 4.**
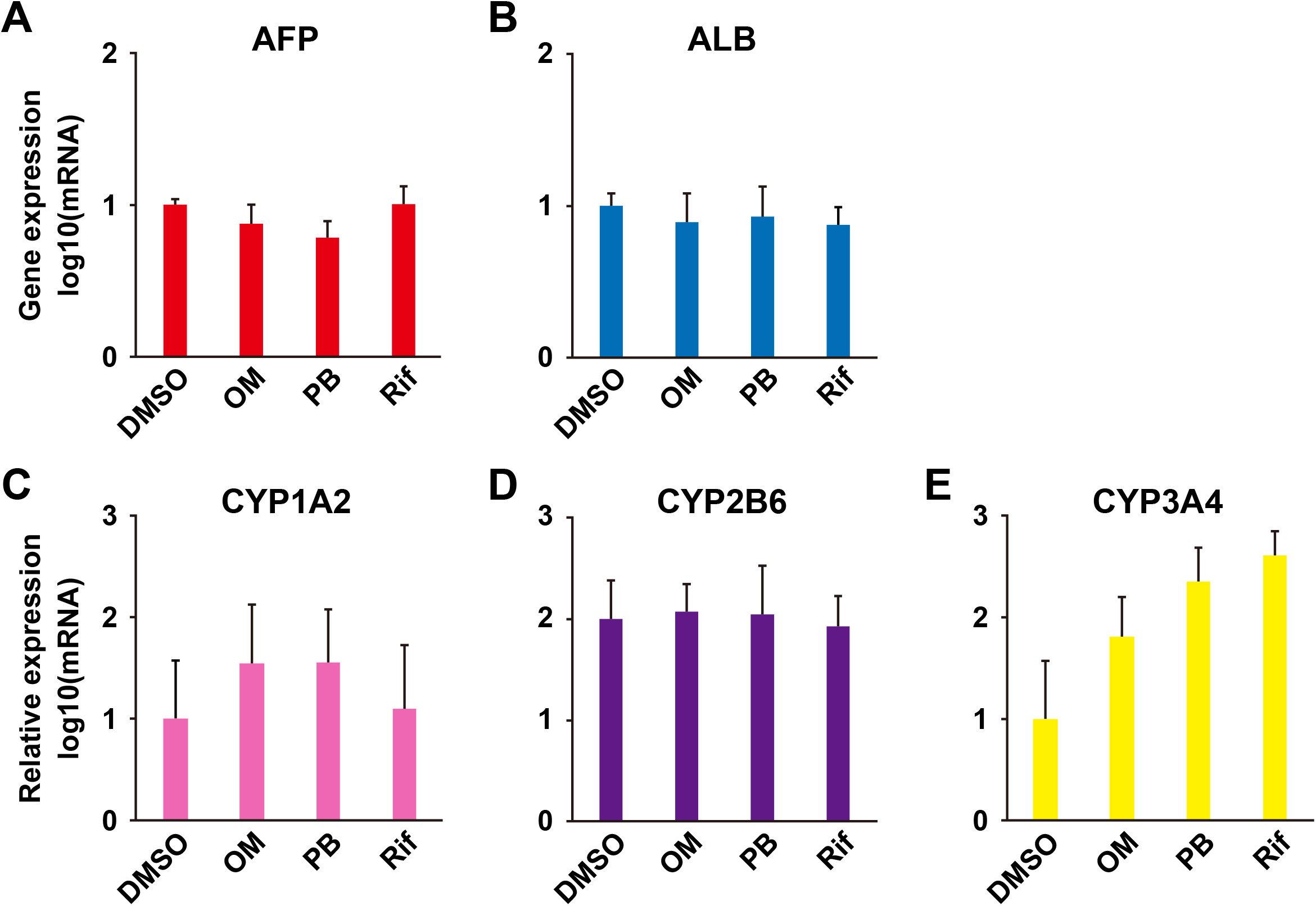
CYP induction with exposure to omeprazole (OM), phenobarbital (PB), and rifampicin (Rif). Expression of AFP (A), ALB (B), CYP1A2 (C), CYP2B6 (D), and CYP3A4 (E) of HepaKI with exposure to DMSO (control), OM, PB, and Rif.

### in vivo analysis of Hepa-KI cells in the kidney

To investigate engraftability and functionality, we implanted human PSC-derived HLCs into the spleen of cDNA-uPA/SCID mice, however, PSC-derived HLCs were not engrafted and only transiently human ALB was elevated. We then implanted organ-derived hepatocytes, i.e. cryopreserved hepatocytes obtained from the surplus liver during liver transplantation in cDNA-uPA/SCID mice. The hepatocytes formed multiple colonies as detected by a human-specific antibody to CYP2C (Figure 5A, B), and blood human ALB was elevated.

**Figure 5.**
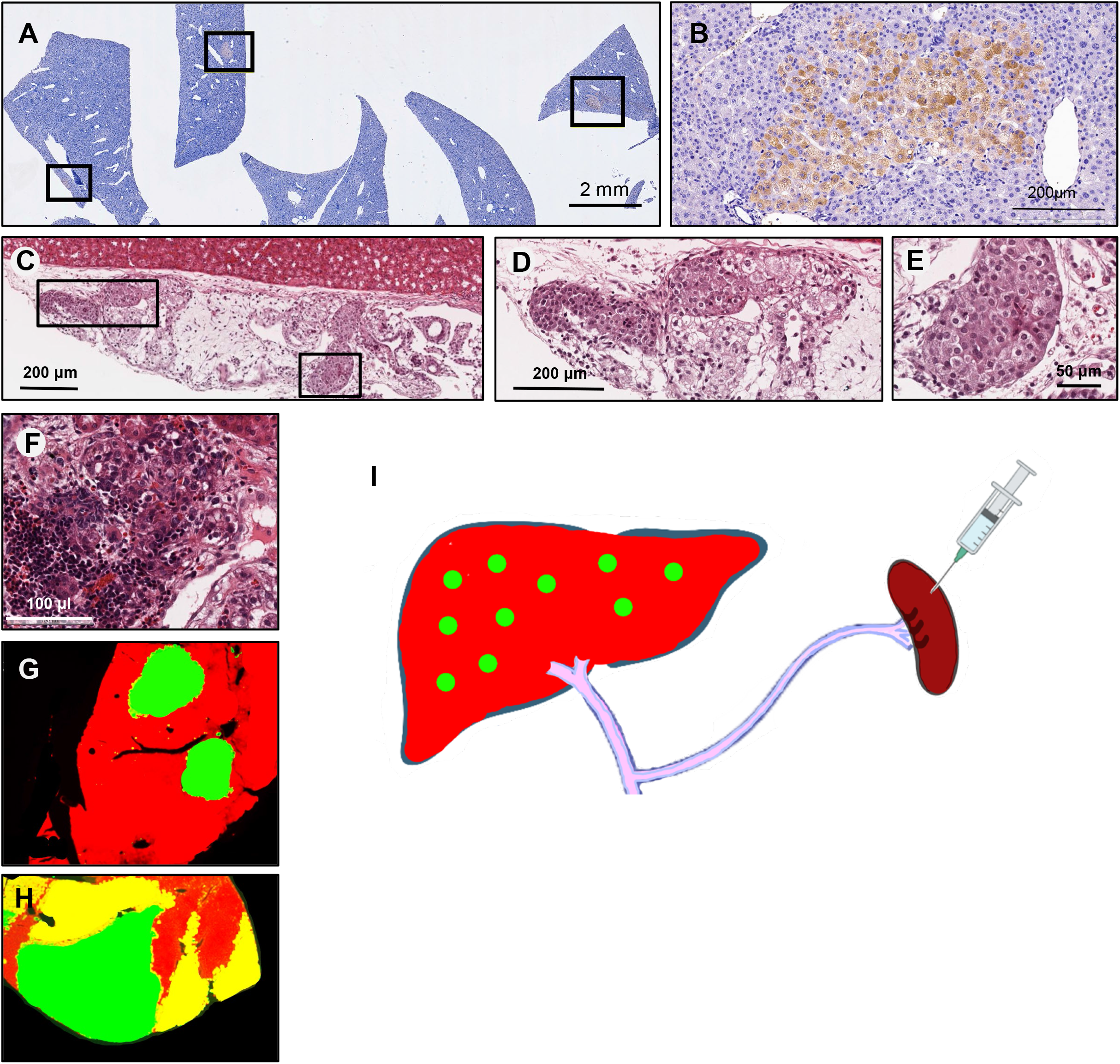
Engraftability of HLCs and hepatocytes. A. Immunohistochemical analysis of hepatocyte colonies in cDNA-uPA/SCID mice, using a human-specific antibody to CYP2C. The hepatocytes formed multiple colonies in the livers 8 weeks after injecting into the cDNA-uPA/SCID mice spleen. Rectangles indicate hepatocyte colonies. B. High-power view of the hepatocyte colony in the liver of cDNA-uPA/SCID mice by immunohistochemistry using the human-specific antibody to CYP2C. C. Implantation of HLCs under the renal capsules of immunodeficient mice. Rectangles indicate hepatocyte colonies. H.E. stain. D. High-power views of the hepatocyte colony under the renal capsules of immunodeficient mice. H.E. stain. E. High-power views of the hepatocyte colony. H.E. stain. F. Extramedullary erythropoiesis along with a hepatocyte colony in the subrenal capsules. H.E. stain. G. Reproduced scheme of double immunofluorescence for human CK8/18 (green) and ornithine transcarbamylase (red) of cDNA-uPA/SCID mice liver that was implanted with hepatocytes from a patient with a hemizygous OTC gene mutation (Figure 1Bc of a paper by Sugahara et al., 2021)^25^. H. Reproduced scheme of double immunofluorescence for human CK8/18 (green) and ornithine transcarbamylase (red) of cDNA-uPA/SCID mice liver that was implanted with hepatocytes from a patient with a heterozygous OTC gene mutation (Figure 1Ba of a paper by Sugahara et al., 2021)^25^. I. Scheme of colony-forming unit in liver (CFU-L). By transplanting hepatocytes into the spleen, hepatocytes form colonies in the liver from the spleen via the splenic vein and portal vein. The number of colonies indicates the number of hepatocytes (hepatic stem cells or hepatic progenitor cells) that can be engrafted and proliferate in vivo. This is the same concept as the formation of colony-forming units in spleen (CFU-s) by hematopoietic stem cells.

We also implanted HepaKI HLCs under the renal capsules of immunodeficient mice. HepaKI successfully engrafted and displayed trabeculae of monomorphic polygonal cells with uniform round/oval nuclei and abundant granular eosinophilic cytoplasm, with capillary vessels (Figure 5C-F). We also observed extramedullary erythropoiesis at the implanted sites in the subrenal capsules (Figure 5F).

## DISCUSSION

In this study, HLCs were prepared from PSCs and characterized as hepatocytes, and engraftment was examined. In vitro, HepaKI showed characteristics as hepatocytes. However, despite the optimization of differentiation protocols, HLCs did not engraft in a mouse model of hepatic failure. There has been a report of engraftment of PSC-derived human HLCs in mice ^32^, and the lack of engraftment in this study may be due to the insufficient protocol for differentiation into hepatocytes that engraft and proliferate in vivo. PSCs themselves have the potential for hepatocyte differentiation, in which hepatocytes clearly appeared as colonies or a cell population in teratomas (Figure 1I, panel for hepatocytes). Distinct colonies of hepatocytes in the teratomas suggest that these colonies are formed from a single hepatocyte/hepatic progenitor/hepatic stem cell during teratoma formation. In addition, HLCs formed trabeculae and exhibited extramedullary erythropoiesis along with colonies of hepatocytes when implanted under the renal capsule (Figure 5D), suggesting a simulation of liver development with the presence of fetal liver cells or hepatoblasts. Hepatocytes in the renal capsule generated a cell population with a trabeculae structure; these trabeculae could have been formed by a large number of differentiated cells or a single proliferative hepatocyte. It is likely that a single hepatocyte proliferates and induces erythropoiesis, similar to that of the embryonic liver.

The CFU-s assay can quantify HSCs in bone marrow, and this idea has contributed to fundamental studies in bone marrow transplantation ^7,8^. Likewise, as for hepatocyte transplantation, we have confirmed that human hepatocytes survived in the liver of immunocompromised mice ^33^. The use of genetically engineered cDNA-uPA/SCID mice, in which liver parenchymal cells are progressively impaired, resulted in more pronounced engraftment of transplanted hepatocytes ^7,8^. The colonies generated by isolated human hepatocytes in the liver of cDNA-uPA/SCID mice are thought to be derived from a single cell. The expression pattern of OTC genes in a single colony is uniform (Figure 5G, H) ^25^, despite the random inactivation of the OTC genes on the × chromosome ^34^. We, therefore, speculate that “colony-forming activity in liver“ can be used to quantify “hepatocytes that engraft and proliferate in vivo“ as a quantitative assay for stem cells that utilize colony-forming ability like bone marrow stem cells (Figure 5I). “Hepatocytes that engraft and proliferate in vivo“ encompass the conceptual notion of liver stem/progenitor cells.

## Acknowledgments

We would like to express our sincere thanks to N. Ito and K. Miyado for the fruitful discussion, to M. Ichinose for providing expert technical assistance, to C. Ketcham for English editing and proofreading, and to E. Suzuki and K. Saito for secretarial work.

## Author Contribution Statement

AU designed experiments. KI, MT, SY, and TY performed experiments. KI and AU analyzed data. KI, MYI, KT, AN, and MK contributed reagents, materials, and analysis tools. MT, HA, HN, TK, and NO discussed the data and manuscript. AU and KI wrote this manuscript.

## Additional Information

The read data have been submitted to the Sequence Read Archive (SRA) under accession number SRP058607.

